# Testing for evolutionary change in restoration: a genomic comparison between *ex situ*, native and commercial seed sources of *Helianthus maximiliani*

**DOI:** 10.1101/2021.03.17.435854

**Authors:** Joseph E Braasch, Lionel N Di Santo, Zach Tarble, Jarrad R Prasifka, Jill A Hamilton

## Abstract

Globally imperiled ecosystems often depend upon collection, propagation, and storage of seed material for use in restoration. However, during the restoration process demographic changes, population bottlenecks, and selection can alter the genetic composition of seed material, with potential impacts for restoration success. The evolutionary outcomes associated with these processes have been demonstrated using theoretical and experimental frameworks, but no studies to date have examined the impact these processes have had on the seed material maintained for conservation and restoration. In this study, we compare genomic variation across seed sources used in conservation and restoration for the perennial prairie plant *Helianthus maximiliani*, a key component of restorations across North American grasslands. We compare individuals sourced from contemporary wild populations, *ex situ* conservation collections, commercially produced restoration material, and two populations selected for agronomic traits. Overall, we observed that *ex situ* and contemporary wild populations exhibited a similar genomic composition, while four of five commercial populations and selected lines were differentiated from each other and other seed source populations. Genomic differences across seed sources could not be explained solely by isolation by distance nor directional selection. We did find evidence of sampling effects for *ex situ* collections, which exhibited significantly increased coancestry relative to commercial populations, suggesting increased relatedness. Interestingly, commercially sourced seed appeared to maintain an increased number of rare alleles relative to *ex situ* and wild contemporary seed sources. However, while commercial seed populations were not genetically depauperate, the genomic distance between wild and commercially produced seed suggests differentiation in the genomic composition could impact restoration success. Our results point towards the importance of genetic monitoring of species used for conservation and restoration as they are expected to be influenced by the evolutionary processes that contribute to divergence during the restoration process.

## Introduction

Restoration aims to mitigate the loss and degradation of native ecosystems by reducing the abundance of non-native species, increasing biodiversity and habitat connectivity, and re-establishing native plant communities resilient to change (Benayas et al. 2009, Hobbs & Norton 1996, Hodgson et al. 2016, Thomson et al. 2009). To achieve these goals extensive inputs of native seed are required, often in quantities too large to be harvested from local, wild populations (Broadhurst et al. 2008, Merritt & Dixon 2013, Pedrini et al. 2020). To compensate for these deficits, seeds used in restoration are often produced commercially to meet increasing demands. However, processes associated with commercial seed production can lead to the evolution of differences that may impact restoration goals (Dyer et al. 2016, Espeland et al. 2017, Nagel et al. 2019, Roundy et al. 1997). Evolution of seed material can occur through a combination of deterministic and stochastic processes, resulting from demographic variation, population bottlenecks, and selection that can influence the amount and type of genetic variation present in restoration material. Bottlenecks and sampling effects following collection, propagation, or cultivation may lead to reductions in genetic diversity and the loss of locally adapted alleles, impacting fitness and reducing the evolutionary potential of restored populations (Blanquart et al. 2013, Fant et al. 2008, Kawecki & Ebert 2004, Robichaux et al. 1997, Williams 2001, Wright 1938). Combined with the impact of selection, which may intentionally or unintentionally lead to genomic and consequent phenotypic change, there is substantial opportunity for evolution of seed material during restoration (Dyer et al. 2016, Espeland et al. 2017, Nagel et al. 2019). Given the impact these different evolutionary processes may have, understanding how these factors interact to influence seed material will have substantial economic and ecological consequences for restoration success (Bischoff et al. 2010, Bucharova et al. 2017, Gerla et al. 2012, Keller et al. 2000, Kimball et al. 2015).

Selection and sampling effects imposed during collection may also pose a significant challenge to the preservation of genetic variation in *ex situ* conservation collections. *Ex situ* seed collections aim to preserve extant genetic variation that may be incorporated into restoration or breeding programs in the future (Hamilton 1994, Li & Pritchard 2009). Both commercial and *ex situ* seed collections aim to maximize genetic diversity while maintaining locally adaptive genetic variation across space and time (DiSanto & Hamilton 2020). Consequently, genomic comparisons between contemporary wild populations, commercially produced material, and *ex situ* conservation collections provide an ideal means to evaluate evolution of seed material maintained for conservation and restoration (Robichauxet al. 1997, Schoen & Brown 1993, Taft et al. 2020). Genomic comparisons of conservation and restoration seed sources with contemporary native populations can be used to infer whether evolutionary challenges inherent to the collection and maintenance of these resources cause them to differ from wild populations they are intended to match.

Sampling effects can generate substantial genomic differences across seed sources with potential lasting impacts to conservation goals and restoration outcomes (DiSanto & Hamilton 2020, Diwan et al. 1995, Franco et al. 2005, Hamilton 1994). The genomic effects of sampling correspond to those found following population bottlenecks, including a reduction in effective population sizes (N_e_) (Leberg 1992, Wright 1938), making this a useful metric to compare across populations when quantifying the effects of sampling. In addition, following a bottleneck, rare alleles are more likely to be lost, influencing the distribution of allele frequencies (Excoffier et al. 2009, Maruyama & Fuerst 1985, Tajima 1989). This loss of rare alleles during bottlenecks results in larger Tajima’s D estimates relative to populations with stable population sizes. Stochastic changes in allele frequencies associated with sampling may also more broadly impact estimation of inbreeding coefficients (F_is_) linked to inbreeding depression (Cavalli-Sforza & Bodmer 1971, García-Cortés et al. 2010, Husband & Schemske 1996) or estimates of coancestry (***θ***) indicative of relatedness among individuals within populations. Importantly, not only are these metrics useful for assessing the magnitude of sampling effects they are also common proxies for evaluating short and long-term fitness due to the challenges associated with inbreeding depression and increased relatedness among breeding individuals (Angeloni et al. 2011, Caballero & Toro 2000, Hughes et al. 2008, Keller & Waller 2002). Thus, these metrics provide valuable comparisons to assess the quality of restoration and conserved seed resources relative to their wild counterparts.

In addition to stochastic processes associated with sampling, directional selection in the agronomic environment can impact seed during cultivation. This may include selection associated with chemical inputs and fertilizers used to improve yield, or reductions in competition or abiotic stress (Dyer et al. 2016, Espeland et al. 2017). Moreover, individuals with traits promoted by mechanized agricultural harvest, such as reduced shattering, minimum heights, or selected phenology could evolve in commercially produced material relative to wild populations (Dyer et al. 2016, Nagel et al. 2019). Previous experimental evidence indicates selection can influence the genetic and phenotypic composition of restoration seed (Dyer et al. 2016, Nagel et al. 2019), but no published studies to date have directly compared the genomic composition of commercially produced seed with native remnant populations in the region of restoration. Genomic signatures of selection can be identified through a variety of statistical analyses developed from the site frequency spectrum (SFS), the distribution of allele frequencies sampled across the genome (Hohenlohe et al. 2010, Nielsen 2001). Of these metrics, Tajima’s D is notable for possessing relatively high statistical power compared to other methods for estimating the strength of selection (Simonsen et al. 1995, Tajima 1983). If selection occurs during commercial seed production, then we would expect commercial populations to have more negative values of Tajima’s D relative to wild populations. Selection and sampling effects in response to the agricultural production are not the only mechanisms that could cause genomic differences between commercial and wild populations. While genetic change may be attributable to anthropogenic selection, natural variation in gene flow may also contribute to genetic differentiation among populations. Isolation by distance (IBD) can create genetic differences among populations and is expected to increase with increasing spatial distance (Slatkin 1993, Wright 1943). Consequently, the relationship between geographic and genetic distance can provide a valuable null hypothesis against which alternative evolutionary scenarios may be tested (Bradburd et al. 2016). If genomic differentiation among seed source populations can be solely explained by the geographic distances between populations, we can conclude that there has not been additional evolution associated with seed source type. However, if IBD is absent or insufficient to explain population differences, other evolutionary factors likely contribute to differentiation across seed source types.

North American grasslands remain one of the most threatened ecosystems globally, with over 98% of remaining habitat converted due to anthropogenic development (Comer et al. 2018, Hoekstra et al. 2004, Samson et al. 2004). However, substantial efforts are ongoing to mitigate the loss of our native grasslands by applying commercially produced restoration seed mixes to reduce non-native species, enhance biodiversity, and increase connectivity across fragmented landscapes (Benayas et al. 2009, Hobbs & Norton 1996, Thomson et al. 2009). The perennial forb *Helianthus maximiliani* Schrad. (or Maximilian sunflower) is a common constituent of grassland communities. *H. maximiliani* encompasses an extensive range of climatic variation, with a natural distribution spanning a broad latitudinal range from northern Mexico to southern Canada (Kawakami et al. 2011, USDA 2004). Previous genetic studies using microsatellites revealed substantial heterozygosity and low inbreeding rates within populations, consistent with an obligate outcrossing mating system (Kawakami et al. 2011). Quantitative genetic experiments have also found differentiation in traits important to adaptation associated with climatic variation, including freezing tolerance and flowering time (Kawakami et al. 2011, Tetreault et al. 2016). There have also been efforts to breed *H. maximiliani* (hereafter selected lines) as a source of seed oil by selecting for increased height, reduced shattering, and increased seed yield (Asselin et al. 2020). *H. maximiliani* functions as a common and effective component of grassland restoration seed mixes due to its ability to readily establish from seed, rapid spread through rhizomatous growth, and ability to reinforce soil structure in degraded habitats (McKenna et al. 2019, USDA 2004). Here, we take a genomics approach to evaluate the factors contributing to evolutionary change among *H. maximiliani* seed sources to inform both conservation and restoration efforts into the future.

Specifically, we compare the genomic composition of wild contemporary, *ex situ*, commercially produced, and agronomically selected seed source populations to (1) test for differences in the genomic composition of different seed source populations using ordination and metrics of genetic differentiation, (2) test whether isolation by distance or alternative evolutionary hypotheses are required to explain genomic differences among seed source populations, and (3) compare population genetic summary statistics across seed sources to indicate potential impacts of sampling and selection. With this third objective, we compare statistics that indicate how much genetic variation is maintained across seed sources as a metric of evolutionary potential, including expected heterozygosity (H_e_), inbreeding coefficients (F_is_), and linkage disequilibrium effective population size (LD-N_e_). We also estimate and compare parameters that can be used to evaluate whether sampling effects or the impact of selection contribute to genomic differences across seed sources. This includes F_is_ and LD-N_e_, in addition to coancestry (***θ***), and Tajima’s D. Overall, this study serves as an important test of recent hypotheses that identify the important role evolutionary processes can play throughout the collection, propagation, and implementation stages of conservation and restoration. Our results provide valuable guidance to both future collection and deployment of seed for restoration and identify new avenues of research that can address the evolutionary consequences seed collection and cultivation have to conservation and restoration.

## Methods

### Population Sampling

We compared the genomic architecture of four distinct seed source types; contemporary collections from wild populations, seed collected and/or cultivated by commercial suppliers for restoration, seed preserved in *ex situ* collections, and lines selected for agronomic traits (hereafter selected lines) to assess the impact demographic variation and unintentional selection may have had on the evolution of seed material used in restoration.

During the summer of 2016, tissue was sampled across six naturally formed wild contemporary populations of *H. maximiliani*, separated by at least 15km, across North Dakota and Minnesota (Figure 1). Leaf tissue was sampled by randomly collecting leaves from 20 individuals per population along a 100m transect (Table 1). Following collection, leaves were preserved in silica gel prior to DNA extraction. Four commercial restoration seed suppliers within North and South Dakota provided five seed populations of *H. maximiliani* for use in this study. Commercial seed was produced either through direct harvest from the wild or by cultivating local genotypes (Table 1). All commercial seeds were harvested between 2016 and 2019.

**Figure 1.**
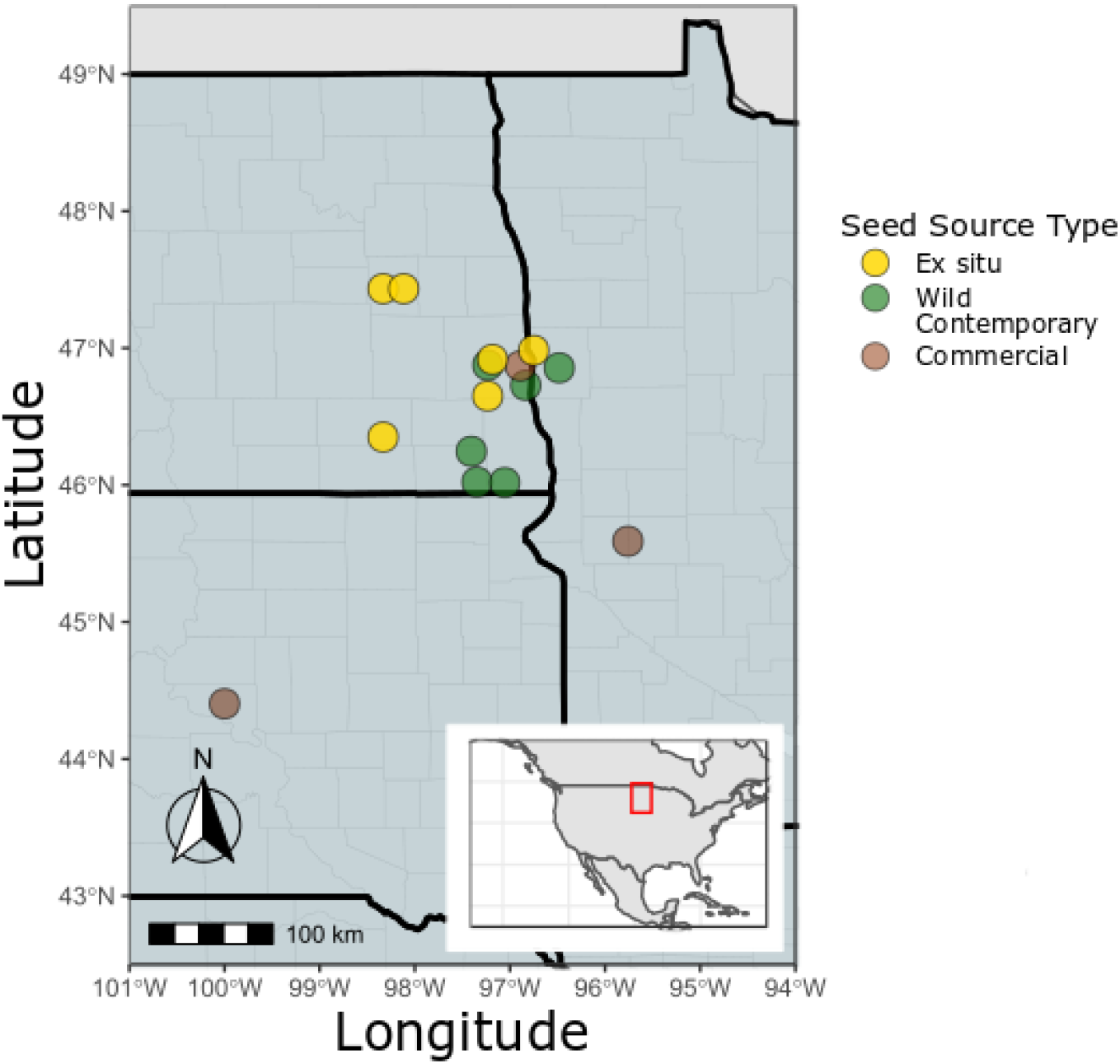
Sampling locations of *Helianthus maximiliani* seed across the northern United States. Location data was available for all native remnant and *ex situ* seed sources and three of the five commercial seed sources used in this study. Location data for the remaining commercially produced seed was not available.

**Table 1.** Geographic location, sample size, and year of harvest for *Helianthus maximiliani* seed sources.

| Seed Source | Population ID | n | State Collected From | Latitude | Longitude | Cultivated | Year of harvest |
| --- | --- | --- | --- | --- | --- | --- | --- |
| *Ex situ* |  |  |  |  |  |  |  |
|  | ES-A | 20 | ND | 47.4333 | -98.3333 | N | 1991 |
|  | ES-B | 20 | ND | 46.9167 | -97.1833 | N | 1991 |
|  | ES-C | 19 | ND | 47.4333 | -98.1167 | N | 1991 |
|  | ES-D | 19 | ND | 46.3500 | -98.3333 | N | 1991 |
|  | ES-E | 20 | MN | 46.9833 | -96.7500 | N | 1995 |
|  | ES-F | 20 | ND | 46.6500 | -97.2333 | N | 1995 |
| *Wild Contemporary* |  |  |  |  |  |  |  |
|  | W-1 | 13 | ND | 46.8758 | -97.2321 | N | 2016 |
|  | W-2 | 20 | MN | 46.8554 | -96.4814 | N | 2016 |
|  | W-3 | 13 | ND | 46.7278 | -96.8339 | N | 2016 |
|  | W-4 | 20 | ND | 46.2459 | -97.4060 | N | 2016 |
|  | W-5 | 20 | ND | 46.0216 | -97.3462 | N | 2016 |
|  | W-6 | 20 | ND | 46.0177 | -97.0537 | N | 2016 |
| *Commercial* |  |  |  |  |  |  |  |
|  | C-1 | 20 | MN | 45.5895 | -95.7600 | N | 2017 |
|  | C-2 | 20 | ND | 46.8697 | -96.8903 | Y | 2018 |
|  | C-3 | 20 | SD | 44.4031 | -99.9997 | Y | 2016 |
|  | C-4 | 20 | SD | - | - | Y | 2016 |
|  | C-5 | 19 | ND | - | - | N | 2017 |
| *Selected Lines* |  |  |  |  |  |  |  |
|  | S-1 | 20 | KS | - | - | Y | 2014 |
|  | S-2 | 20 | KS | - | - | Y | 2014 |

Note:
n: number of individuals in each population retained for genetic analysis.
State Collected From: MN, Minnesota; ND, North Dakota; SD, South Dakota.
Cultivated: N, seed collected from naturally growing stands; Y, grown in an agroecosystem for at least one generation prior to seed harvest

*Ex situ* seed populations included in this study were sourced from the USDA National Genetic Resources Program (https://www.ars-grin.gov/). These bulk seed collections, designated by local provenance, were collected in North Dakota in September of 1991 (4 collections) and 1995 (2 collections). *Ex situ* seeds were bulk harvested by clipping mature seed heads, following which seed heads were dried and cleaned prior to storage in a cold room at 4 ℃ with 25% humidity.

Selected lines represent germplasm developed as part of a domestication program to improve the agronomic value of perennial grassland species. Breeding populations of *H. maximiliani* were originally founded with 10 plants from each of 96 wild populations (960 plants total) harvested in Kansas, US. Each line was bred for five generations selecting for increased yield per stem, yield per seed head, and seed size by pooling pollen from the twenty best performing families in each generation and using pooled pollen to fertilize plants from the same twenty families. Seeds were harvested in 2014 and stored at 4 ℃.

To obtain leaf tissue for genomic analysis, in 2018 we grew seeds sourced from commercial, *ex situ*, and selected lines indoors and under growth chamber conditions in Fargo, ND. Seeds were germinated in bulk following the protocol by Seiler (2010). Achenes were surface-sterilized, soaked for 15 minutes in a 2% solution of 5.25% sodium hypochlorite in distilled water with a single drop of wetting agent (Tween 20, Sigma-Aldrich, Inc. St. Louis, MO, USA). Achenes were then rinsed and scarified with a razor blade, cutting through the hull and tip of cotyledons before soaking in a 100 PPM solution of gibberellic acid (Sigma-Aldrich, Inc.) for 60 minutes. Following this, achenes were placed onto filter paper in Petri plates, sealed with parafilm, and stored overnight in the dark at 21℃. Seeds (embryos) were gently removed from hulls, returned to Petri plates, and examined daily for germination. Seeds with a visible radicle were planted into a moistened peat pellet (Jiffy Peat Pellet, Plantation Products, Norton, MA, USA) and grown at 21℃ under artificial lights (fluorescent T12 bulbs) until they produced between 4-8 true leaves. Leaf tissue samples were collected from 20 randomly selected individuals from each population or seed collection (20 individuals x 13 sources = 260 total individuals) and stored in silica gel prior to DNA extraction.

### DNA sequencing and genotyping

We extracted DNA from ~10mg of dried leaf tissue using a modified Macherey-Nagel NucleoSpin Plant 2 extraction kit that included additional ethanol washes to ensure removal of secondary plant compounds. DNA concentration was verified using the Quant-iT™ PicoGreen^®^ dsDNA kit (Life Technologies, Grand Island, NY) after submission to the University of Wisconsin-Madison Biotechnology Center for sequencing. Genomic libraries were prepared as in Elshire et al (2011) with minimal modification. In short, 50 ng of DNA was digested using the 5-bp cutter ApeKI (New England Biolabs, Ipswich, MA) after which barcoded adapters were added by ligation with T4 ligase (New England Biolabs, Ipswich, MA) for Illumina sequencing. The 96 adapter-ligated samples were pooled and amplified to provide library quantities appropriate for sequencing, and adapter dimers were removed by SPRI bead purification. Fragment length and quantity of DNA was measured using the Agilent Bioanalyzer High Sensitivity Chip (Agilent Technologies, Inc., Santa Clara, CA) and Qubit^®^ dsDNA HS Assay Kit (Life Technologies, Grand Island, NY), respectively. Libraries were standardized to 2nM and were sequenced using single read, 100bp sequencing and HiSeq SBS Kit v4 (50 Cycle) (Illumina Inc.) on an Illumina HiSeq2500. Cluster generation was performed using HiSeq SR Cluster Kit v3 cBot kits (Illumina Inc, San Diego, CA, USA).

Sequence files were demultiplexed using *ipyrad* version 0.9.12 (Eaton 2014) allowing for zero mismatches in the barcode region. Following demultiplexing, single nucleotide polymorphisms (SNPs) were called across populations and seed sources using the dDocent v2.7.8 pipeline (Puritz et al 2014a, b). In the first step of the pipeline, reads were trimmed using the program TRIMMOMATIC (Bolger et al. 2014), including the removal of low-quality bases and Illumina adapters. Following read trimming, the pipeline aligned reads to the *Helianthus annuus* v1.0 genome using BWA (Li & Durbin 2009). Sequence alignment was performed using the software’s default parameters (a match score of 1, mismatch score of 4, and gap score of 6). Finally, as a last step, dDocent called SNPs using the software FREEBAYES (Garrison & Marth 2012) that produced a VCF file with 4,735,557 total SNPs. Downstream SNP filtering of the VCF file first removed missing loci variants with conditional genotype quality (GQ) < 20 and genotype depth < 3. Then, loci with Phred-scores (QUAL) ≤30, allele counts < 3, minor allele frequencies < 0.05, call rates across all individuals < 0.9, mean depth across samples > 154 (based on the equation from Li et al. 2014), and with linkage scores > 0.5 within a 10kb window were removed. Following downstream filtering, 12,943 polymorphic loci were kept and used for subsequent analyses. Individuals with more than 30% missing genotypes were removed from the analysis. In total, 14 individuals from wild contemporary populations, two individuals from *ex situ* collections, and one individual from a commercial supplier were discarded, leaving a total of 363 genotyped individuals for inclusion in subsequent analysis (Table 1).

### Population structure and genetic differences between seed sources

To test for the effects of seed source on the genetic structure among *H. maximilani* populations, we used principal components analysis (PCA) and discriminant analysis of principal components (DAPC) to partition the genetic variation observed in our sampling. Pairing these methods provides valuable insight as it allows comparison of a method agnostic to *a priori* expectations for population structure (PCA) to one which attempts to best depict differences between populations (DAPC). Additionally, while PCA allows for the visualization of individual axes which explain decreasing amounts of the total genomic variation, DAPC can combine and display variation across multiple axes of variation simultaneously. Thus, DAPC will isolate and incorporate only those axes that contribute to differences between our populations, while PCA depicts population groupings onto major axes of variation.

We performed principal components analysis (PCA) on the matrix of SNPs used for all individuals in the study. Missing data (2.5% of all loci) were substituted with the mean allele frequencies at each locus. We calculated the total variation explained by each axis by dividing the eigenvalue of each PCA and the total sum of all eigenvalues. PCA was performed with the *dudi.pca* function within the ADEGENET package (Jombart 2008, Jombart & Ahmed 2011). We then plotted individuals along the first two PCA axes using *s.class* function in the package ADEGRAPHICS (Siberchicot et al. 2017).

We first applied DAPC to the entire SNP dataset and then to a subset of the data including only individuals from wild contemporary and *ex situ* populations. DAPC partitions genetic variance using principal components analysis before using discriminant analysis to maximize interpopulation variation and minimize intrapopulation variation. This allows DAPC to identify the axes of variation that simultaneously maximize between group differences and minimize within group differences (Jombart et al. 2010). For both analyses, we retained principal component axes sufficient to explain 90% of the total variation and retained 18 and 11 discriminant functions for depicting between group differences for all seed sources and *ex situ –* wild contemporary analysis, respectively. All DAPC analyses were performed using the R package ADEGENET (Jombart 2008).

### Isolation by distance in seed collections

Genomic differences across populations can arise from the independent evolution of populations connected by limited gene flow giving rise to isolation by distance (IBD). Patterns of neutral evolution produced by IBD could confound our ability to infer evolutionary changes caused by selection associated with seed source type. For this reason, we tested for any correlation between F_st_ calculated between two populations and the spatial distance between them. Testing for IBD was also necessary due to the uneven spatial distribution of populations from different seed sources to confidently attribute genomic differences to environmental or sampling effects associated with different seed source types.

Pairwise genetic differences between populations were calculated as F_st_ using the Weir and Cockerham’s method which is unbiased with regard to differences in sample sizes (Weir & Cockerham 1984; Willing et al. 2012). Unlike DAPC, pairwise F_st_ allows us to quantify the total genetic differences between population pairs. Importantly, we can compare the magnitude of genetic differences for populations of the same seed source type (e.g. two wild contemporary populations) to differences calculated between populations of different seed source types (e.g. wild contemporary population versus commercial population). If evolved differences between populations have developed due to conditions associated with seed source type, we expect inter-source F_st_ to be larger than intra-source F_st_. To test for differences in inter- and intra-source Fst, we used a Wilcoxon rank sum test implemented with the function ‘wilcox.test’ in R.

To test for effects of IBD and seed source types on pairwise F_st_, we first calculated the geographic distance between seed collections. Exact geographic location data was available for all wild contemporary and *ex situ* populations. Locations for commercial populations C-1, C-2, and C-3 were estimated as approximate locations based on descriptions of the counties, cities, reserves, and geographic features associated with the provenance for each collection. Provenance data was not available for the remaining two commercial populations (C-4, C-5) and selected lines (S-1, S-2), and therefore these populations were not included in the analysis. Geographic distances were calculated using the haversine formula, which accounts for the curvature of the earth (Robusto 1957), and then square root transformed to improve model fits. Distance measurements were made using with the R package GEODIST.

The relationship between F_st_, distance, and seed source types used to calculate F_st_ was evaluated using model selection with a series of linear mixed models. In these models F_st_ was expressed as function of spatial distance (fixed effect) and a factorial variable coded for the different pairwise seed source comparisons with random slopes and intercepts. The variable for seed source comparisons required six levels in total, three for each of the intra-source comparisons (wild to wild, *ex situ* to *ex situ*, and commercial to commercial) and three for each of the inter-source comparisons (wild to *ex situ,* wild to commercial, and *ex situ* to commercial). We compared the full model which included both spatial distance and seed sources to two reduced models each of which included only one of the terms. A likelihood ratio test, implemented with *lrtest* function in the package LMTEST (Zeileis & Hothorn 2002), was used to identify significant differences between the full and reduced models. When the full and reduced models were significantly different, we chose the model with the greatest loglikelihood value as the model with the best fit.

Additional calculations of per-locus F_st_ supported earlier analyses of population structure between seed sources, particularly for comparisons including selected lines.

### Signatures of Sampling and Selection

To ascertain the importance of sampling effect and selection in contributing to differences among seed sources, we calculated expected heterozygosity (H_e_), inbreeding coefficients (F_is_), linkage disequilibrium effective population size (LD-N_e_), and coancestry coefficients (***θ***). Expected heterozygosity (H_e_) and inbreeding coefficients (F_is_) were calculated individually for each SNP using the R package ADEGENET v2.1.0 (Jombart 2008, Jombart & Ahmed 2011). To estimate LD-N_e_, we followed the method of Braasch et al. (2019) which, rather than producing a single, genome wide value, uses the mean from a distribution of LD-N_e_ estimated using multiple subsets of the data. This method reduces the likelihood of violating the assumption of no physical linkage among loci in organisms without assembled genomes by repeatedly sampling a smaller subset of loci. To produce a distribution of LD-N_e_ estimates, we created 5,000 sets of 500 loci and estimated LD-N_e_ for each using the function ldNe in the package StrataG (Archer et al. 2017) following methods from Waples et al. (2016). Estimates of relatedness (***θ***) were made using the R package COANCESTRY (Wang 2011) with 2,000 bootstrap iterations to calculate 95% confidence intervals for each population.

We compared H_e_ and F_is_ across seed source types using linear mixed models with seed source type as a fixed effect and population as a random effect. The significance of individual terms and post hoc tests were performed with the R package LMERTEST (Kuznetsova et al. 2017). Differences between LD-N_e_ and ***θ*** among wild contemporary, *ex situ,* and commercial seed sources were compared using linear models implemented with the *lm* function. Selected populations were not included in linear models due to lack of replication (n=2).

To test for signatures of selection or bottlenecks across seed source types, we calculated Tajima’s D for each population (Tajima 1989). Positive estimates of Tajima’s D are indicative of high heterozygosity or a scarcity of rare alleles, which could be caused by balancing selection or demographic bottlenecks, as well as sampling effects. Conversely, recent population expansion or directional selection should result in negative values of D arising from an excess of rare alleles. Whole genome estimates of Tajima’s D with accompanying p-values were produced with the ‘tajima.test’ function in the R package PEGAS and compared across seed source types with an analysis of variance (ANOVA) and Tukey post hoc test (Paradis 2010).

We also note here that plotting per-locus F_st_ as a function of H_e_ revealed that the data were depauperate in low H_e_, high F_st_ loci. This pattern has been found in other work and matches the expected relationship for these variables when drift and selection contribute similarly to evolutionary differentiation (Narum & Hess 2011). When drift and selection similarly impact the genome, accounting for neutral differentiation in outlier analysis could increase type II error while failing to account for it would increase type I error. As a result, outlier analyses are not expected to yield reliable results and were therefore not considered in this manuscript.

We found commercial populations had greater values of Tajima’s D than wild contemporary populations (see results). This difference could be caused by either the loss of rare alleles or greater genetic diversity in commercial populations. To visualize the frequency of rare alleles and overall genetic diversity across seed source types, we constructed a folded site frequency spectrum (SFS) for each seed source, with the exception of the selected lines. SFSs were estimated from the filtered SNPs dataset (12,943 variants, 363 individuals) using the set of R functions available at https://github.com/shenglin-liu/vcf2sfs. Individuals from populations classified as the same seed source type (Table 1) were pooled together to generate seed source-specific allele frequency profiles (Figure. 3).

## Results

### Population structure and genetic differences between seed sources

In total, 363 individuals and 12,943 SNP loci passed our filtering requirements. PCA of the entire dataset required 110 axes to explain over 50% of the total genetic variation across all seed source types. A total of 4.0% of the total genetic variation was explained by the first principal component axis, which differentiated the two selected populations from all other seed sources (Figure 2A). The second axis explained 2.2% of the total genetic variation and separated *ex situ* population ES- E and commercial populations C-2 and C-5 on either end of the axis and from all remaining populations at the center.

**Figure 2.**
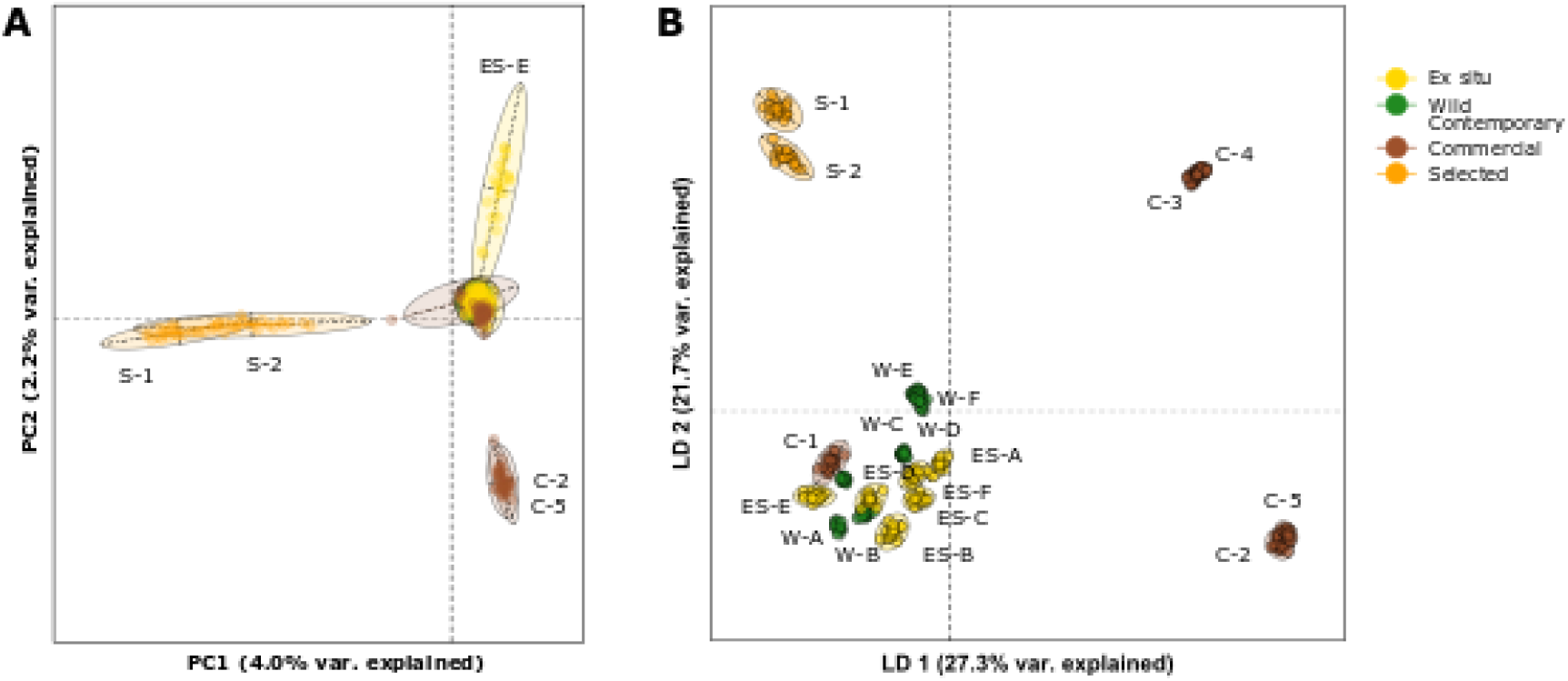
Genomic variation of *Helianthus maximiliani* partitioned by (A) principal components analysis (PCA) and (B) discriminant analysis of principal components. Both analyses were conducted on the full SNP dataset for all *ex situ*, wild contemporary, commercial, and selected populations. Missing data (2.5% of all observations) in the PCA were replaced with the mean allele frequency value. Different seed source types are depicted as different colors (yellow: *ex situ*; green: wild contemporary; brown: commercial; orange: selected).

**Figure 3.**
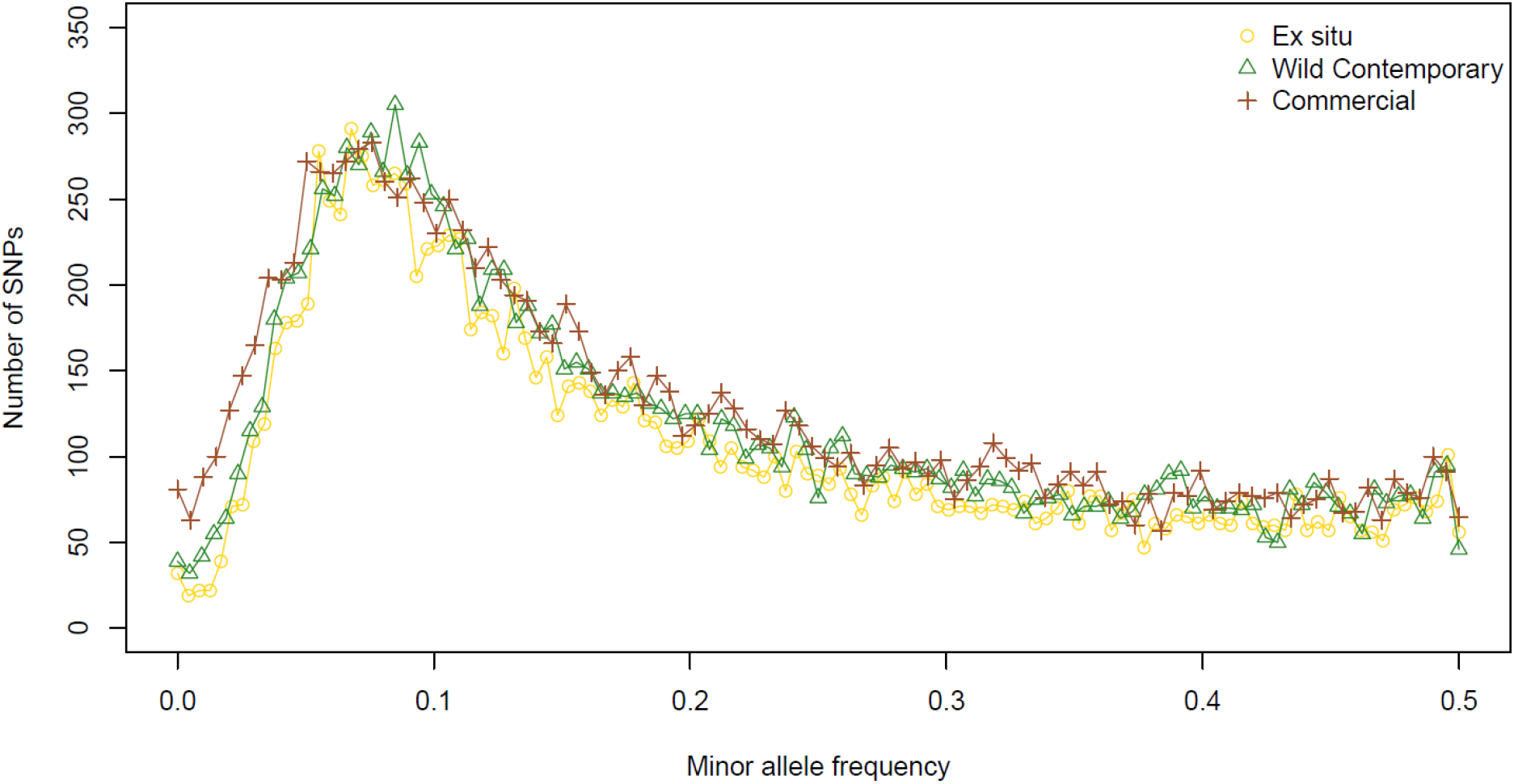
The site frequency spectrum for wild contemporary, *ex situ*, and commercial seed collections shows a greater number of low frequency alleles in commercial populations.

When genetic variation was partitioned using DAPC the first two axes explained 27.3% and 21.7% of genetic variation, respectively (Figure 2B). These values are considerably greater than PCA-axes because DAPC incorporates and depicts the relationships across multiple axes of variation simultaneously. DAPC, which also attempts to maximize differences between pre-defined groups, split populations of *H. maximiliani* into four distinct groups. All wild contemporary, all *ex situ*, and commercial population C-1 from Minnesota formed one group together. The remaining four commercial populations were split into two clusters according to the state they were sourced from. Commercial populations from North Dakota, C-2 and C-5, grouped together, as did the commercial populations from South Dakota, populations C-3 and C-4. Selected populations formed their own unique cluster. The first DAPC axis split commercial populations, except for commercial population C-1, from wild, *ex situ*, and selected populations. The second DAPC axis split wild and *ex situ* genotypes from selected genotypes and split the North Dakota and South Dakota commercial populations.

A DAPC including only *ex situ* and wild contemporary seed explained 47.6% of the total variation in this subset of the data (Figure S1). The first and second ordination axes explained 32.3% and 15.3% of genomic differences, respectively. Populations did not split according to seed source type, although most *ex situ* populations were on the left side of axis 1 and the bottom of axis 2. Each population formed a distinct cluster, except for two *ex situ* populations (ES-B and ES-E) which grouped together.

### Isolation by distance in seed collections

Pairwise F_st_ ranged from −0.001, between commercial populations C-2 and C-5, to 0.238 between *ex situ* populations ES-E and selected population S-1 (Figure S2). Pairwise F_st_ mirrored patterns observed in PCA. The largest values of F_st_ observed were between selected and non-selected populations. Additionally, F_st_ values calculated with population ES-E were larger than F_st_ calculated using any other *ex situ* collections. Across all pairwise F_st_, inter-seed source comparisons were significantly greater than intra-seed source comparisons (Wilcoxon signed rank test: W = 3845, p < 0.001) (Figure S3). The linear mixed model using pairwise F_st_ as the dependent variable and seed source comparison as the only independent variable was significantly better than the full model which included seed source comparison and geographic distance (Table 2). The model using only distance as an independent variable was not significantly different from the full model.

**Table 2.** The effect of isolation by distance and seed source on pairwise F_st_. Reduced models were compared to the full model using the likelihood ratio test. P-values in bold indicate significant differences between the reduced and full model.

| Model | Log-likelihood | Df | P-value |
| --- | --- | --- | --- |
| Distance + Seed Source Comparison | 223.75 | - | - |
| Distance | 223.49 | 1 | 0.42 |
| Seed Source Comparison | 228.98 | 1 | **0.001** |

### Patterns of genetic diversity and relatedness

Average H_e_ for all *H. maximiliani* populations ranged between 0.211±0.002 SE and 0.275 ±0.001 SE, while F_is_ estimates ranged from −0.019 ±0.001 SE to 0.018 ±0.002 SE (Figure 4A, C). H_e_ was similar across wild contemporary, *ex situ*, and commercial populations, all of which were significantly greater than H_e_ in selected lines (Full model: F_3,15_ = 6.9, P = 0.004) (Posthoc tests: wild contemporary-selected: P<0.001, *ex situ*-selected: P=0.001, commercial-selected: P=0.002) (Figure 4A). There were no significant differences in F_*is*_ across seed source types (F_3,15_ = 1.1, P = 0.363).

**Figure 4.**
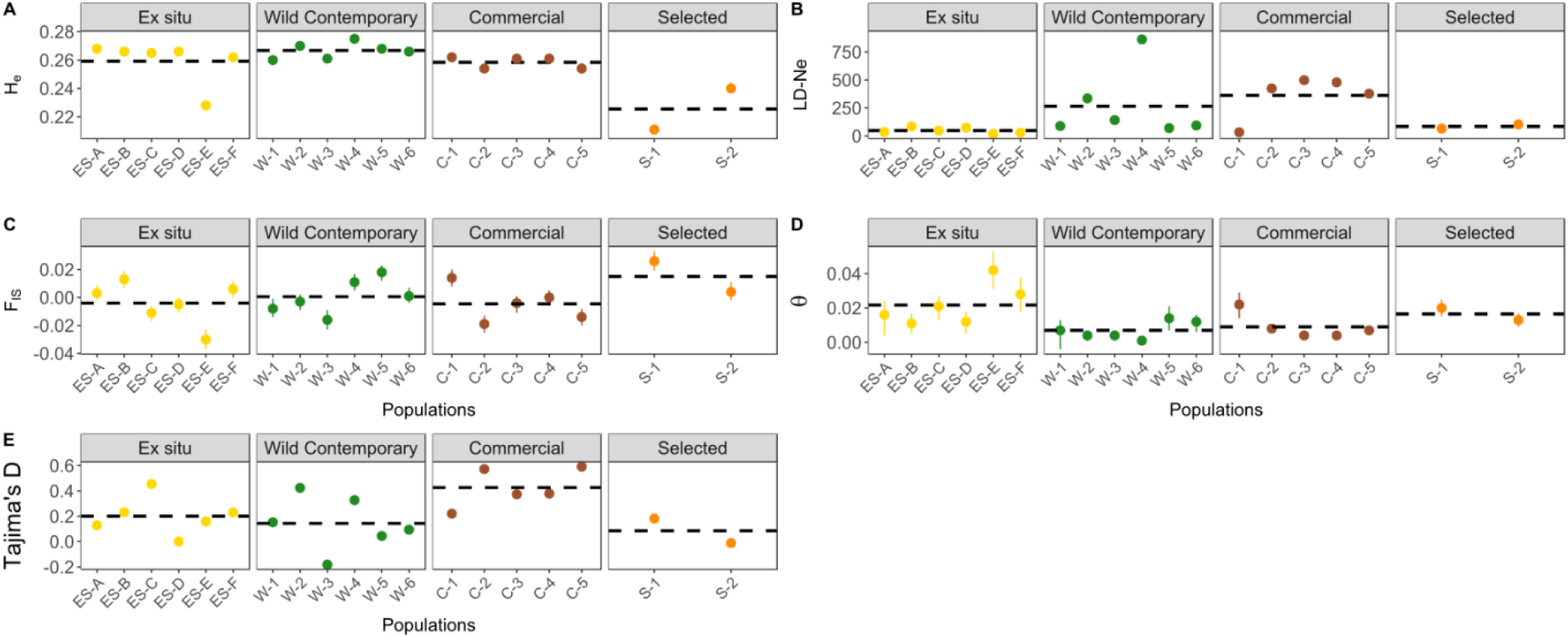
Estimates of (A) expected heterozygosity (H_e_), (B) linkage disequilibrium effective population size (LD-N_e_),(C) inbreeding coefficients (F_is_), (D) coancestry coefficients (θ), and (E) genome wide Tajima’s D for commercially produced, *ex situ*, native remnant, and experimentally selected *Helianthus maximiliani* seed sources. Points depict population means for panels A-D. Error bars for LD-N_e_ and θ are bootstrapped 95% confidence intervals estimated with 2000 replicates.

**Figure 5.**
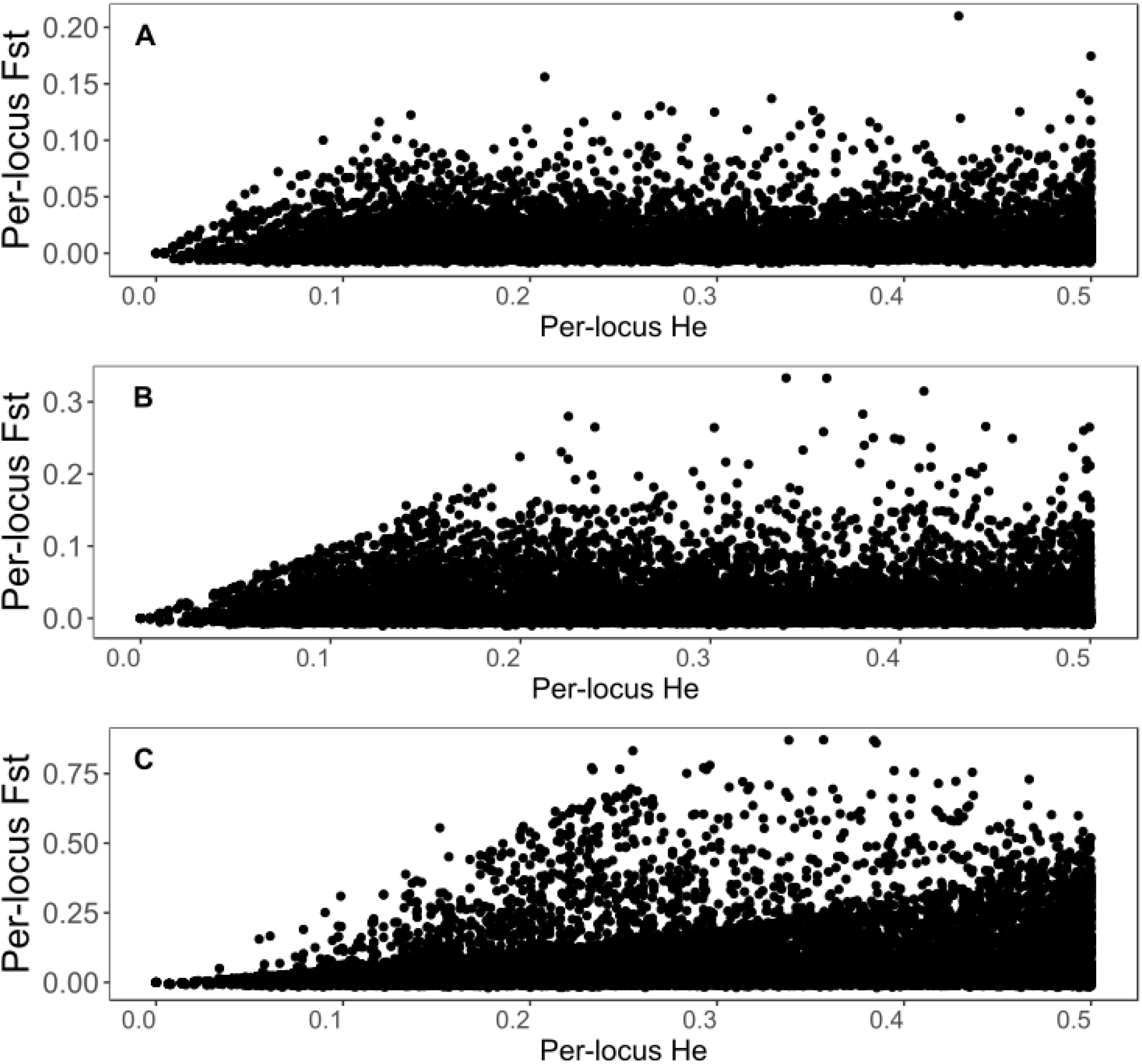
The per-locus relationship between F_st_ and He for a) wild and *ex situ* populations, b) wild and commercial populations, and c) wild and selected populations. In all comparisons, there are no loci with low He and moderate to high F_st_, which matches patterns from simulation studies in which the effects of selection and neutral evolution are equivalent in magnitude.

Wild contemporary and commercial seed populations spanned a wide range of LD-N_e_ estimates in comparison to *ex situ* and selected populations (Figure 4B). Despite these trends we did not observe significant differences in LD-N_e_ between seed source types (F_2,14_=3.3, P = 0.069). Patterns of coancestry across seed source types mirrored those for LD-N_e_. The linear model comparing the effect of seed source was significant (F_2,14_=3.6, P = 0.037). Although wild contemporary and commercial populations had lower ***θ*** (range 0.001 to 0.022) than *ex situ* and selected populations (0.011 to 0.042), post hoc comparison revealed only e*x situ* and commercial populations were significantly different.

Genomewide estimates of Tajima’s D were not significantly different from neutral expectations for any population, including selected lines (Table S1). Nonetheless, Tajima’s D estimated for commercial populations was significantly greater than those for wild contemporary populations (analysis of variance: F_2,14_ = 3.82, P = 0.048; Tukey post hoc test wild – commercial: P = 0.047) suggesting differences in the site frequency spectrum among populations for these two seed sources. The shape of the folded SFS for wild, *ex situ*, and commercial seed sources was similar, with few rare alleles, a peak at a frequency around 0.09, and a gradual decline at higher frequencies. The SFS for commercial genotypes could be distinguished from *ex situ* and wild genotypes by a higher abundance of the rarest alleles.

## Discussion

An overarching goal of both restoration and conservation is to maintain evolutionary potential to ensure populations sustain the ability to adapt to change (Hamilton et al. 2020, Hoffmann & Sgro 2011). However, for both *ex situ* conservation collections or seed propagated for restoration, the efficacy of these goals may be dependent upon the amount and type of genetic variation maintained in populations. Sampling effects and genetic bottlenecks associated seed collection and selection during propagation can create genotypic differences between seed source types. Using a genomic dataset assembled from wild contemporary, commercial, *ex situ,* and selected populations of *H. maximiliani*, we tested for the presence of genomic differences that could be attributed to seed source type. We found evidence that commercial seed and selected lines were genetically differentiated from wild and *ex situ* collections. These differences could not be explained by neutral processes, such as isolation by distance, implicating other evolutionary explanations for genomic differences among seed sources. While we did not find direct evidence that selection caused genomic differentiation between seed sources, increased coancestry and low LD-N_e_ in *ex situ* collections were consistent with an impact of sampling. Varying genomic composition of commercial seed sources relative to wild, contemporary populations suggest further study is required to evaluate whether genomic differences correspond to functional differences that impact restoration success. Common garden studies have shown that seed transfer across environments can impact plant traits and performance (Bucharova et al. 2017, Giencke et al. 2018, Johnson et al. 2004, Lesica & Allendorf 1999, Yoko et al. 2020). Consequently, the genomic differences we observe here warrant additional study of *H. maximiliani* seed sources linking genomic differences to traits important to adaptation and persistence in restored environments.

Consistent with the expectation that genotypic differences exist between seed source types, we observed genetic differences between *H. maximiliani* populations according to seed source type and region of origin. While *ex situ* and wild populations appear to have similar genomic composition, with the exception of one *ex situ* population, commercial and selected populations tended to exhibit differences across our analyses. Differences between seed source types were most apparent in the DAPC analysis. Although this method maximizes between group differences and minimizes within group differences, our analysis grouped individuals according to population, rather than seed source type. Seed source differences were also apparent with PCA, which split selected lines from all others along the first axis of variation. PCA also grouped all remaining populations, except for commercial populations C-2 and C-5, which had mostly negative values along the second PCA axis, and *ex situ* collection ES-E, which had more positive values along the second axis. Pairwise comparisons of F_st_ matched ordination analyses, and in general F_st_ calculated between collections of the same seed source type were lower than those calculated across seed source types. Overall, the differences we observed between wild and commercial seed match the expectations established by theoretical and experimental studies for how evolution via selection and sampling effects could lead to differentiation between commercial and wild populations (Dyer et al. 2016, Espeland et al. 2017, Nagel et al. 2019).

Unexpectedly, DAPC also split the four divergent commercial populations according to the state of their collection, either North or South Dakota. The split among commercial populations was also apparent with PCA, which separated the two commercial populations from North Dakota from all other non-selected populations. Notably, within an individual region, commercial populations were grown by different suppliers despite their genomic similarity. Genetic similarity among different sources of commercial seed could indicate consistent local practices in commercial production or that suppliers are pulling from similar genetic resources. We did not find robust evidence for selection as a cause for the differences between wild and commercial seed, which suggests it is more likely that seed suppliers are using similar genetic stock. Regardless of the reason for this effect, similarity in commercial seed does not match most conservation goals which attempt to balance high genetic diversity with the need for locally adapted seed inputs (Hamilton et al. 2020, Hufford & Mazer 2003, McKay et al. 2005), a problem compounded when commercial seed is not a close analogue to wild populations. Commercial seed is used for restoration because the necessary volume of seed cannot be sustainably harvested from wild populations (Broadhurst et al. 2008). If there are few *H. maximiliani* populations of appropriate size for harvesting seed within different regions, it would then be unsurprising if different commercial suppliers obtained and mixed germplasm from the same wild sources. While we don’t have information on how well the seed used in this study compares to wild genotypes as a restoration resource, the dissimilarity between commercial and wild seed warrants greater communication between seed suppliers and restoration practitioners to understand the potential causes of genomic differences.

Genomic differences between *H. maximiliani* populations were not correlated with geographic distances and do not appear to demonstrate patterns of IBD. In natural populations, genomic differences are expected to increase in response to increasing spatial distance and a corresponding reduction in gene flow among populations (Slatkin 1993, Wright 1943). The absence of IBD in our data could have multiple explanations. First, there could be sufficient gene flow to connect *H. maximiliani* populations across the largest spatial scales included in our analysis, but substantial gene flow should also homogenize the genomic variation between populations. This does not correspond to the results of our DAPC analysis, which was able to partition genomic variation, not just at the scale of seed source types, but at the level of individual populations. An alternative cause for the lack of IBD could be rates of gene flow near zero, such that every population is functionally isolated, negating the effect of distance. Although fragmentation of prairie habitat in North America has indeed increased the isolation among plant populations (Samson et al. 2004; Wimberly et al. 2018), the complete cessation of gene flow across populations has not been observed in other species. In the grass *Festuca hallii*, distance was still correlated with genetic variation across the same geographic region considered in our study (Qiu et al. 2009). Although grasses and sunflowers differ in their pollination ecology and methods of seed dispersal, these patterns of differentiation in *F. hallii* suggest it is unlikely that prairie plant populations are so isolated that geographic distance has no effect on population structure. Rather, given the structure of our analysis, it is more probable that seed source differences disrupted patterns of IBD and more strongly predicted differences in pairwise F_st_. Increased sampling across commercial and wild populations would be useful to supplement our observations on the effect of seed source type and warrants additional study. Knowledge of the degree to which commercial propagation disrupts the effects of natural IBD in *H. maximiliani* populations would be valuable for restoration practitioners seeking to best match seed inputs to local environmental conditions.

Selection during agricultural propagation can result in the evolution of restoration seed, altering traits that contribute to growth and phenology (Dyer et al. 2016, Nagel et al. 2019). Although commercial populations were genetically distinct from wild contemporary populations, we did not find evidence that differences are due to selection. Commercial and wild populations did not differ in H_e_, F_is_, LD-N_e_, or coancestry. Tajima’s D in commercial populations was also not significantly different from zero, which suggests that selection has not been strong enough to exert genome scale effects. Interestingly, Tajima’s D was significantly greater in commercial than wild populations, which is likely caused by a slight increase in the frequency of rare alleles in commercial populations. Although selection in agricultural ecosystems is common, experimental cultivation of five different plant species found that molecular evidence of evolution was not apparent in two (Nagel et al. 2019). Species that were perennial or outcrossing, such as *H. maximiliani,* were also less likely to exhibit evidence of selection. Thus, although we uncovered multiple ways in which commercial and wild populations differ, the life history and mating system of *H. maximiliani* may have buffered the species against evolutionary change during commercial production. Overall, the genomic differences between commercial and wild populations do not appear to be driven by selection during cultivation, a phenomenon which might be more common in plant species with shorter life histories or that exhibit greater instances of selfing.

We found significant differences in coancestry between *ex situ* and commercial seed sources. *Ex situ* populations also had lower LD- N_e_ than commercial populations, and although this comparison was not significant, a high coancestry should coincide with higher rates of linkage disequilibrium and lower LD-N_e_. Low LD-N_e_ and higher coancestry without corresponding increases in F_is_ could reflect the sampling methods used to establish these collections. Alleles are more likely to be identical by descent in populations with greater coancestry and are less likely to represent the uniform sampling of large populations (Cavalli-Sforza & Bodmer 1971). In *ex situ* collections, high coancestry and low LD-N_e_ could result from sampling large quantities of seed from a relatively small number of maternal individuals. Sampling in this manner would also not immediately reduce H_e_ or increase F_is_ in a self-incompatible species prior to sexual reproduction (Allendorf 1986, Leberg 1992), but would increase coancestry and LD-N_e_ because of the large number of half-siblings represented in the population. The difference between commercial and *ex situ* collections may imply that commercial seed provides a superior resource by harboring greater genotypic diversity. Whether or not this is true likely depends on the specific goal of the collection. For example, high coancestry could be mitigated if multiple *ex situ* collections are mated prior to deployment in the wild. Additionally, *ex situ* collections appear to be closer analogues to contemporary wild populations and could be superior resources for restoration if the genotypic differences depicted in our analysis correlate with functional differences. This suggests additional work to evaluate the consequences of high coancestry and genomic differences from wild populations will be essential for applying our results into practice for restoration.

The production of seed for restoration and conservation includes an inherent conflict between maintaining the genomic composition of wild populations and supplying large volumes of seed (Broadhurst et al. 2008, Espeland et al. 2017). In addition to these challenges, the goals of conservation are themselves sometimes in conflict, with the need to maintain populations that are locally adapted while maximizing genetic diversity to buffer against contemporary and future environmental challenges respectively (Bucharova et al. 2017, Hamilton et al. 2020). The loss of genetic diversity and evolution of functional traits during cultivation is thus a major concern for restoration efforts. In our comparison of commercial and wild *H. maximiliani* collections, we did not find evidence of selection or reduced genetic variation in commercial seed, but we did observe significant differences in their genotypic composition. Additionally, the surprising genomic similarity of commercial seed sourced from the same region is evidence for a homogenizing factor either during seed collection or cultivation. High similarity across commercial seed inputs is at odds with the goal of maximizing genetic diversity while maintaining local adaptation and has the potential to reduce the efficacy of restoration in the short and long-term. Given the species-specific evolutionary consequences of cultivation (Nagel et al. 2019), it is also possible that other seed inputs which are less buffered against the genomic effects of selection, due to their life history or mating strategies, will exhibit increased differentiation from wild populations during commercial production. Additional study evaluating the trait variation and contribution of *H. maximiliani* to ecosystem services between wild and commercial seed collected across varied restored habitats is necessary. Furthermore, to fully integrate the consequences of our study for restoration similar work comparing plant species commonly used in restoration will be important for generalizing these results. Until this work can be performed, increased collaboration between producers and users of commercial seed is needed to better understand the effects of provenance, individual methods of harvest, and cultivation on seed material needed to best meet restoration goals (Hamilton et al. 2020).

## Acknowledgments

We would like to thank the commercial seed producers who provided seed material for use in this study. In addition, we would like to thank Anna Bucharová, Malte Conrady, Jess Lindstrom, Dan Syvertson, and Kate Volk, along with X anonymous reviewers for suggestions which greatly helped improve this paper. We also would like to thank Joshua Miller, Katie Lotterhos, and Rhiannon Peery for assistance with our analyses. This work was supported by a USDA-NACA (#433203), and a new faculty award from the office of the North Dakota Experimental Program to Stimulate Competitive Research (ND-EPSCoR NSF-IIA-1355466) to JAH, funding from NSF-RESEARCH-PGR-1856450 to JEB, and funding from the NDSU Environmental and Conservation Sciences Program to LND.

