## Supplemental Tables and Figures for "Testing for evolutionary change in restoration: a genomic comparison between *ex situ*, native and commercial seed sources of *Helianthus maximiliani*"

Table S1. Tajima’s D values for all populations. P-vales indicate whether Tajima’s D values deviate from neutral expectations.

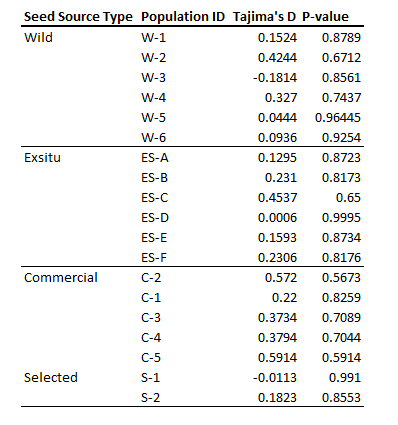

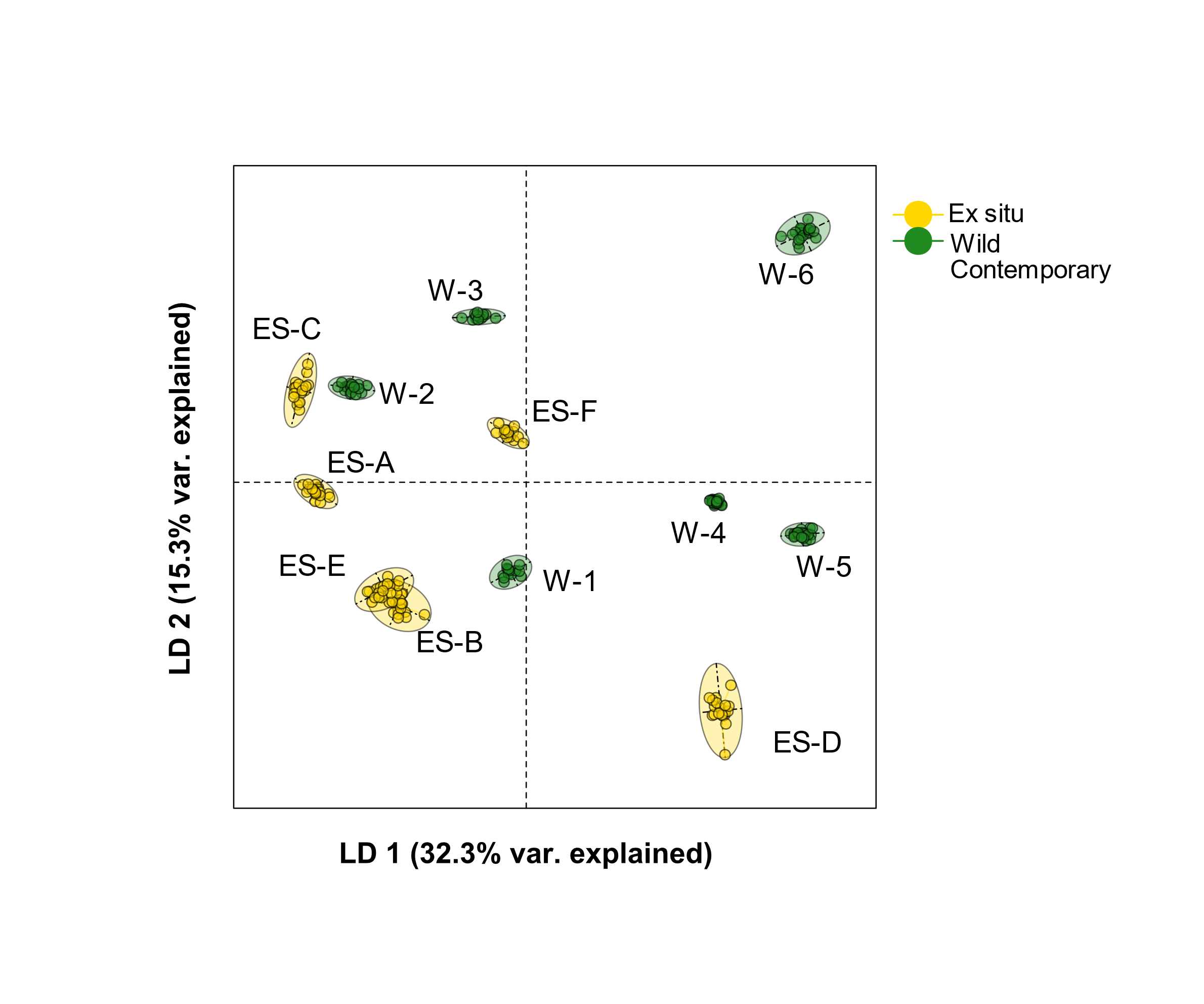

Figure S1. Discriminant analysis of principal components for *Helianthus maximiliani* SNP genotypes from *ex situ* and wild contemporary populations only. Different seed source types are depicted as different colors (yellow: *ex situ*; green: wild contemporary).

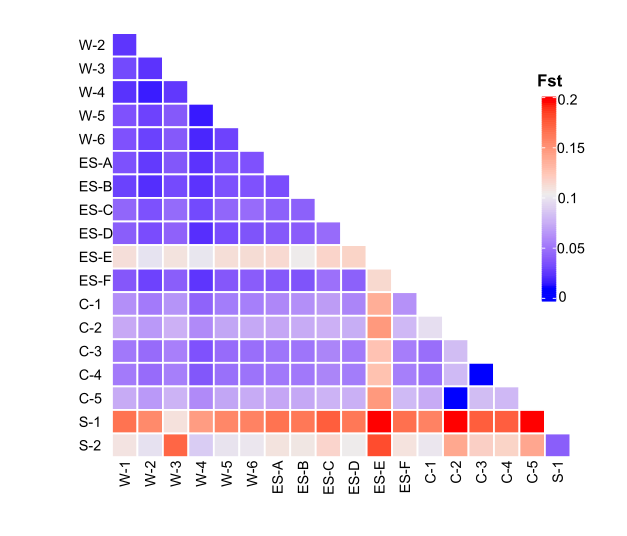

Figure S2. Heatmap of pairwise F_st_ for 19 populations of *H. maximiliani*. F_st_ was calculated using the Wier and Cockerham method.

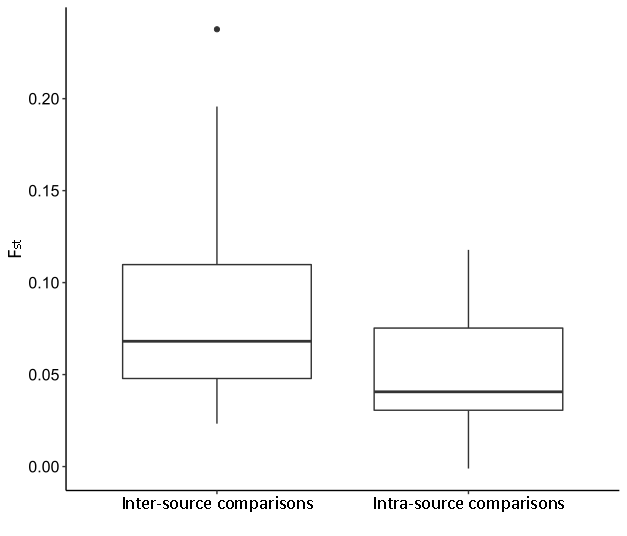

Figure S3. Comparisons between pairwise-Fst calculated for intra-seed source comparisons (both populations are the same seed source type) are lower than inter-population comparisons (the two populations are different seed source types) (P < 0.001).
